# Lupeol, a bioactive compound of tamarind, functions as a potent inhibitor of hepatitis C virus (HCV) entry through disruption of the E2–CD81 interaction

**DOI:** 10.64898/2026.07.22.739987

**Authors:** Maitri Singh, Chayan Bhattacharjee, Avik Bardhan, Aparna Mukhopadhyay

**Affiliations:** Department of Life Sciences, Presidency University, Kolkata-700073, West Bengal, India

**Author notes:** Both authors contributed equally (Maitri Singh, Chayan Bhattacharjee). Corresponding Author: Dr. Aparna Mukhopadhyay, Department of Life Sciences, Presidency University, 86/1 College Street, Kolkata=700073. CB is currently located in Premas Biotech, Gurugram, Haryana, India. AB is currently located in IISER Pune, India. This work is part of two theses submitted/ to be submitted in partial fulfilment of a PhD degree from Presidency University, Kolkata.

**Keywords:** Hepatitis C virus (HCV), Phytochemicals Antiviral agents, HCV entry inhibition, HCV pseudoparticles (HCVpp), Molecular docking, ADME screening, Toxicity endpoints, Target prediction, Molecular dynamics (MD) simulation, Lupeol, Natural product-based therapeutics, Homology modeling, Drug discovery

## Abstract

Hepatitis C virus (HCV) infection remains a major global health challenge despite the success of direct-acting antivirals (DAAs), which are limited by high cost, restricted accessibility, and the emergence of resistant strains. Natural products, particularly phytochemicals, represent a promising reservoir of antiviral agents with diverse mechanisms of action and favorable safety profiles. In this study, we combined wet-lab experimentation with computational approaches to identify plant-derived molecules capable of inhibiting HCV entry. Guided by ethnobotanical evidence, methanolic leaf extracts of Psidium guajava L., Plumeria alba L., Syzygium cumini L., and Tamarindus indica L. were prepared and evaluated for cytotoxicity in Huh7 hepatoma cells. Entry inhibition was assessed using EGFP-labelled HCV pseudoparticles (HCVpp) by qRT-PCR and confocal microscopy. Among the tested plants, Tamarindus indica extract significantly reduced KGFP expression (p < 0.05), confirmed by ΔΔCq analysis and impaired membrane fusion events, while Psidium guajava and Plumeria alba impaired intracellular trafficking without blocking initial attachment. Syzygium cumini showed no inhibitory effect under the tested conditions. Complementary in silico analyses included homology modelling, molecular docking, ADME/toxicity profiling, and molecular dynamics simulations of HCV E2–ligand complexes. Literature mining identified 39 candidate compounds, among which lupeol exhibited stable binding interactions with HCV E2 and favorable pharmacokinetic properties. Critically, in vitro binding assays confirmed that lupeol disrupted the E2–CD81 interaction, reducing bound E2-EGFP to 6% compared to controls. This was supported by HCV-pseudoparticle entry assays confirming inhibition of entry. Together, these findings establish Tamarindus indica and lupeol as potent HCV entry inhibitors.

## 1. Introduction

Hepatitis C virus (HCV) infection remains a major global health challenge, with more than 71 million people chronically infected worldwide^1^. As a blood borne, enveloped, positive-sense RNA virus, HCV establishes lifelong infection that progressively damages the liver, driving the transition from fibrosis to cirrhosis and sometimes hepatocellular carcinoma^2^. This progression accounts for a considerable share of hepatitis-related morbidity and mortality^2^. Viral entry into hepatocytes is mediated by the envelope glycoproteins E1 and E2, which form a heavily glycosylated heterodimer^3^. The E2 glycoprotein binds to the large extracellular loop of the tetraspanin receptor CD81, a critical step in viral attachment and entry^3^. Disrupting this E2–CD81 interaction strongly reduces HCV entry in cell models, making it a promising target for entry inhibitors^4^.

Direct-acting antivirals (DAAs) have transformed HCV therapy, offering high cure rates^5^. Yet, they remain costly, show variable efficacy across genotypes, and do not prevent reinfection^5^. Access is limited in low-resource settings, and host-targeting agents raise safety concerns^6^. These limitations underscore the urgent need for complementary strategies^6^. Natural products, particularly phytochemicals derived from medicinal plants, represent a compelling alternative^7^. They are locally available, cost-effective, and rich in diverse bioactive compounds with fewer side effects. Flavonoids, tannins, and triterpenoids have already demonstrated inhibitory effects against HCV replication and entry, highlighting their potential as novel antiviral leads^7, 8^.

Different plant parts—including roots, bark, flowers, and seeds—are known to contain phytochemicals with antimicrobial and antiviral activity^8^. However, leaves are the most abundant and easily accessible plant material, making them practical for collection in sufficient quantities. Their richness in secondary metabolites further supports their use in phytochemical and antiviral studies. Guided by ethnobotanical knowledge and literature, we selected four locally abundant species from Kolkata—*Psidium guajava* (guava), *Plumeria alba* (champa), *Syzygium cumini* (jamun), and *Tamarindus indica* (tamarind)—all known for hepatoprotective and antiviral properties (Table 1). These plants have long history of use in traditional Indian medicine and contain compounds such as gallic acid and quercetin with reported antiviral activity.

**Table 1.**
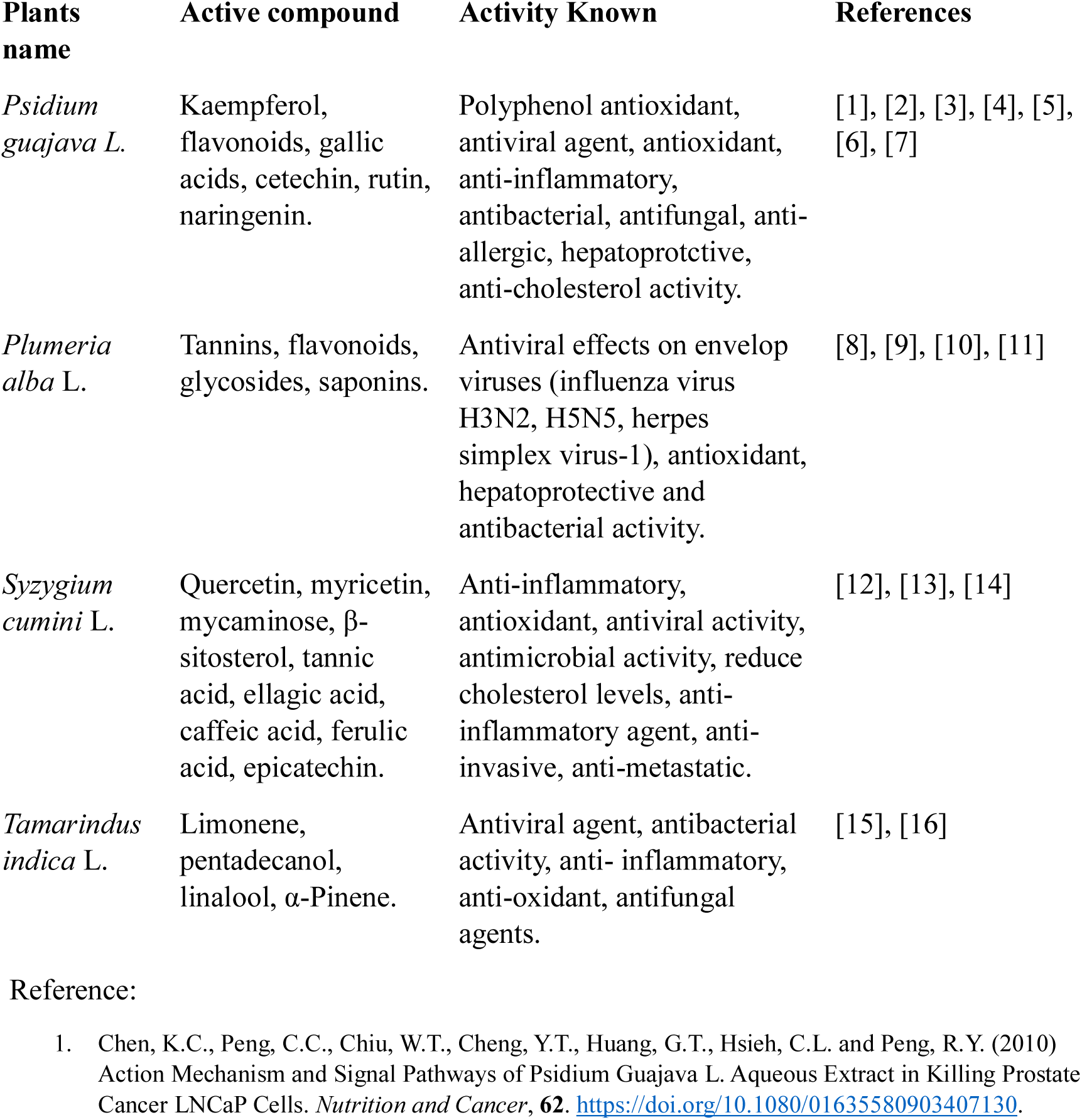
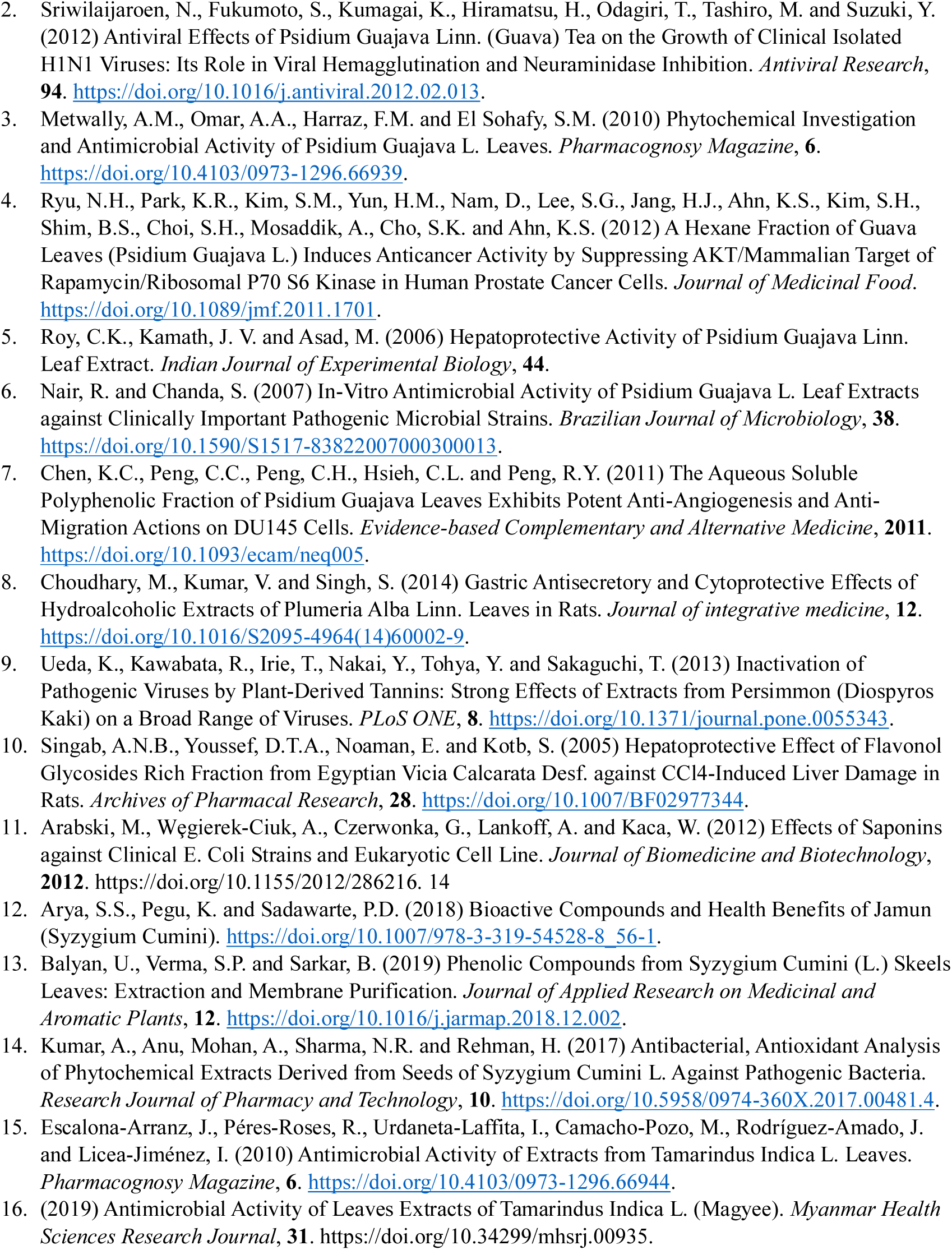
Compounds found in plant leaves and their pharmacological effect.

In this study, we integrated literature-guided phytochemical selection with computational approaches including homology modelling, molecular docking, ADME/toxicity profiling, and molecular dynamics simulations. In parallel, methanolic leaf extracts were tested in Huh7 cells using EGFP-labelled HCVpp to assess entry inhibition. This combined strategy identified plant-derived inhibitors of the conserved E2–CD81 interaction, offering affordable, entry-targeted antiviral leads that could complement DAAs. This is particularly relevant to patients requiring frequent blood transfusions, which is a major source of contracting new infections.

## 2. Materials and Methods

### 2.1. Literature Screening and collection of plant samples

Through a literature survey, four ethnobotanically important plants available in Kolkata were selected. Leaf samples of *Psidium guajava* L. (guava), *Plumeria alba* L. (white champa), *Syzygium cumini* L. (black plum), and *Tamarindus indica* L. (tamarind) were collected between March and April from different regions of Kolkata and authenticated by Dr. Debabrata Maity, Assistant Professor, Department of Botany, University of Calcutta. Healthy leaves were air-dried for seven days at room temperature, powdered using a grinder, and subjected to solvent-gradient extraction.

### 2.2. Preparation of leaf extracts

Leaf extracts of *Psidium guajava*, *Plumeria alba*, *Syzygium cumini*, and *Tamarindus indica* were prepared by sequential cold extraction using solvents of increasing polarity (n-hexane, chloroform, methanol, water). For each sample, ∼50 g powdered leaf was extracted with 200 mL of solvent at room temperature (24 °C) with agitation at 120 rpm for 24 h; fractions were decanted after each step and the residue re-extracted with the next solvent. Extractions were performed with assistance from Prof. Binay Chaubey. Collected fractions were concentrated by rotary evaporation, reconstituted in DMSO to 100 mg/mL, sterile-filtered through 0.22 μm membranes, and stored at −20 °C.

### 2.3. Cell viability assay

Cell viability and cytotoxicity of all methanolic leaf extracts were assessed in Huh7 cells using the methylene blue assay across a concentration range of 10 µg/mL to 1000 µg/mL, following a published protocol^9^. Based on the preliminary data using methylene blue (data not shown), only concentrations more than 50 μg/ml were tested by MTT assay following the protocol of Pandey et al^5^. In brief, 2×10^4^ Huh7 cells/well was plated into 96-well plates (Nunclon Delta Surface, Thermo Fisher Scientific, Denmark). Various concentrations (50 to 1000μg/ml) of methanolic leaf extracts were added to the cells and incubated at 37°C for 48h. DMSO was used as control. Post-treatment, the culture medium was aspirated from each well, and 10 µL of MTT solution (3-(4,5-dimethylthiazol-2-yl)-2,5-diphenyl tetrazolium bromide, Sisco Research Laboratories Pvt. Ltd. (SRL) India) was added. 3 hours post incubation media was carefully removed and 100μl of DMSO was added to dissolve the formazan crystals and luminescence measured with a microplate reader (Synergy/ H1, Biotek, USA) at 540 nm and a reference wavelength of 620nm.

### 2.4. Preparation of fluorescent labelled HCVpp, plasmid isolation and purification

All experiments in this study were conducted in either Human embryonic kidney cells HEK293T for cellular transfection, Human hepatoma cell line Huh7 for viral transduction experiments^10^. HEK 293T cells were grown in DMEM media, Huh7 cells in RPMI media, containing L-glutamine (Gibco, Thermofisher Scientific, USA), supplemented with 10% FBS and 1% penicillin streptomycin. pcDNA-E1E2 plasmid containing full length of HCV (genotype 1a) E1E2 and retroviral structural gene gag-pol containing construct pTG5349, were obtained as a kind gift Prof. Jean Dubuisson, Université Ede Lille, France. GFP reporter containing *Lentiviral* construct pLenti-PGK-GFP was obtained from Dr. Jayanta Roychoudhury, Albert Einstein College of Medicine, NY, USA. EGFP tagged construct was generated from pEGFP-N1 plasmid (Clontech)^11^. Fluorescent labelled HCVpp was generated using our standardized protocol^11^. In short, three constructs encoding pEGFP-E1E2, PTG5349/gag-pol and pLenti-PGK-GFP were simultaneously transfected into HEK293T cells using calcium-phosphate method (Calphos Mammalian Transfection kit, Clontech as per manufacturer’s instructions). 40h post transfection HCVpp containing media was collected purified using 0.45µm filter (Millipore, MILLEX-GV) and concentrated using Amicon ultra filter (Amicon Ultra Centrifugal Filters, Merck). Concentrated HCVpps were stored at -80C till future usage.

### 2.5. Studies of HCV entry inhibition with methanolic leaf extracts

Huh7 cells were either grown on sterile coverslips or on 60×15mm cell culture dishes. 50µl HCVpp along with methanolic plant extracts (Supplementary Table 1) were added drop wise to the cells. After 30min of incubation on ice, cells were washed with cold PBS and HCVpps allowed to internalise by incubation at 37°C for 20min. HCVpp transduced Huh7 cells were thereafter shifted to ice to block further entry. Entry of HCVpp was tested via two methods. Either the presence of KGFP was determined by quantitative real-time PCR (qRT-PCR) or localisation of E1E2GFP was observed via microscopy. The following primer sets were used to determine the presence of KGFP (Forward-5’-CATTCTGCAAGCCTCCGGAG-3’ and Reverse-5’-CTTGTACAGCTCGTCCATGC 3’). GAPDH estimation served as loading control, estimated using Forward-5’-GCACCGTCAAGGCTGAGAAC-3’ and Reverse 5’-TGGTGAAGACGCCAGTGGA-3’). The fold change of KGFP expression for all the transduced cells was determined by 2^ ^-ΔΔ Cq^ calculation to yield absolute values. Briefly, ΔCq values were obtained by subtracting GAPDH Cq from KGFP Cq in each sample. ΔΔCq was then calculated by subtracting the ΔCq of Huh7 cells transduced with HCVpp only (control) from the ΔCq of Huh7 cells transduced with HCVpp in the presence of plant extracts. Relative fold changes were thus expressed as 2^^-ΔΔCq^, with the control normalized to 1. Values below 1 indicate reduced KGFP expression (inhibition of viral entry), whereas values above 1 indicate relative increase compared to control.

Colocalization of EGFP (E1E2-EGFP) with plasma membrane was determined by a plasma membrane marker FM™ 4-64 (Molecular probes, Invitrogen detection technologies). Images were acquired using confocal microscopy (Leica DMi8) with 63X oil immersion objective. *z*-Series images were acquired in 1024 × 1024 format with scanning speed 100 Hz and sequential imaging manner. Images were pseudocolored and colocalisation measured using ImageJ-Fiji ((1.52a, ImageJ-win-64)) image processing software. Pixel intensities of EGFP or integrated density (IntDen) from all the microscopic images were quantified by Image J (after subtraction of background. IntDen of each of the images for all the time points were normalized with mean IntDen of 0 min and all the replicates (N = 3) were considered.

### 2.6. Homology modelling

The three dimensional structure of HCV1aE2 was obtained from Singh et al, 2025 ^12^.

### 2.7. Selection and preparation of plant derived ligands

A literature survey targeting *Tamarindus indica* L. and HCV identified 39 natural compounds (Supplementary Table 2); searches were performed in PubMed and ScienceDirect using keywords such as “HCV,” “Tamarindus indica,” and “natural compounds.”

### 2.8. Preparation of ligands

Three-dimensional (3D) structures of the selected ligands were retrieved from the PubChem (https://pubchem.ncbi.nlmnih.gov/)^13^ and ChemSpider (http://www.chemspider.com/ )^14^ databases. For compounds not available in these repositories, molecular structures were manually constructed using MarvinSketch (version 20.2.0, ChemAxon Ltd., 2020).

### 2.9. Organ toxicity and toxicity endpoints analysis

Toxicity was predicted using the PKCSM (http://biosig.unimelb.edu.au/pkcsm/prediction) web server, which employs graph-based signatures from canonical SMILES to estimate pharmacokinetic and toxicological endpoints^15^. The compound’s SMILES was submitted and retrieved descriptors included molecular weight, AMES mutagenicity, oral rat acute toxicity (LD₅₀), hepatotoxicity, minnow toxicity, hERG-I inhibition potential, and human maximum tolerated dose. PKCSM predictions were used to complement experimental data and guide safety evaluation.

### 2.10. ADME screening

An in-silico ADME assessment was performed using the SWISSADME web server (http://www.swissadme.ch) to evaluate pharmacokinetics, drug-likeness, and medicinal-chemistry friendliness of candidate molecules; analyses included Lipinski’s rule of five to screen for oral bioavailability and suitability for drug development^16^.

### 2.11. Molecular docking and virtual screening

Targeted docking was performed with AutoDock Vina^17^ (Trott and Olson 2010) using a focused grid box covering the defined E2 binding pocket (Supplementary Table 3); protein–ligand contacts were analyzed in BIOVIA Discovery Studio Visualizer, 2019. Virtual screening of 39 compounds (in-house scripts) prioritized ligands that bound favorably across all five HCV E2 subtypes, with selection defined as an average affinity ≤ −8 kcal·mol⁻¹; the top five ligands were advanced for further evaluation.

### 2.12. Target prediction

Target prediction for the hit compound was performed using SwissTargetPrediction web server^18^ (http://www.swisstargetprediction.ch), submitting the canonical SMILES to generate a ranked list of likely macromolecular targets based on structural similarity; this identified potential off-targets, cross-reactivity, and phenotypic liabilities to guide downstream validation.

### 2.13. MD simulation of HCV E2-ligand complexes

Molecular dynamics simulations of the protein–ligand complex were run via the WebGRO server^19^ (GROMACS backend) on fully solvated systems^20^ using default equilibration and production settings; trajectory analyses assessed structural flexibility and stability by calculating RMSD, RMSF, radius of gyration (Rg), hydrogen-bond profiles, solvent-accessible surface area (SASA), and per-residue area to evaluate the complex’s dynamic behaviour.

### 2.14. Selection of ligand for laboratory validation and stock solution preparation

From the initial set of 39 plant-derived compounds, lupeol was selected for subsequent experimental validation. This selection was based on its favorable docking performance, ranking as the second-best candidate in AutoDock Vina analyses, as well as supporting evidence from previously published studies reporting antiviral and antimicrobial activities of lupeol ^21,22,23,24^. Lupeol (Lupeol | 545-47-1 | Tokyo Chemical Industry (India) Pvt. Ltd.) was accurately weighed (1 mg) and dissolved in 1 mL of absolute ethanol to obtain a stock solution with a final concentration of 1 mg/mL. From this stock, working solutions were prepared, and a concentration of 5 µg/mL was used for subsequent experimental assays.

### 2.15. Transient transfection and binding assay

HEK293T cells were seeded in 6-cm culture dishes and grown to approximately 70% confluence prior to transfection. Cells were transfected individually with CD81-LEL-His and pEGFP-E1E2 constructs using Turbofect reagent, according to the manufacturer’s protocol (Invitrogen). For CD81, the media was collected 48 hours post transfection as the protein is secreted. The media was clarified by centrifuged at 1000x g for 10 minutes. For pEGFP-E1E2 transfected cells, cells were lysed in buffer containing 50 mM Tris (pH 7.6), 150 mM NaCl, 0.5% Triton X-100, 5 mM MgCl₂, and 0.5% PMSF. Lysates were clarified by centrifugation at 10,000 × g for 10 min, and the resulting supernatants were used for binding assays. Ni-NTA agarose beads (250 µL) were washed three times with PBS and subsequently incubated with CD81-containing supernatant for 12 h at 4 °C with gentle rotation (22 rpm). Unbound supernatant were retained, and beads were washed to remove non-specific proteins. Binding interactions were assessed by incubating the CD81-bound beads with lysates from pEGFP-E1E2–expressing HEK293T cells. For inhibitor studies, Lupeol dissolved in ethanol was added at a final concentration of 5ug/ml during incubation with the pEGFP-E1E2 lysate. Samples were incubated for 4 h at 4 °C, followed by collection of unbound supernatants and PBS washes of the beads. The presence of E2 was confirmed by immunoblotting using a mouse monoclonal anti-GFP antibody (Thermo Fisher Scientific, MA5-15256).

### 2.16. Statistical analysis

All statistical analyses were performed using SSS 20.0.and Microsoft Excel (Microsoft Excel 2010). Multiple comparisons were conducted by one-way analysis of variance followed by Bonferonni post hoc analysis. Correlation analysis was performed by linear correlation analysis of Pearson’s R (using above threshold values) by Fiji image processing software (ImageJ-win-64). t-score was determined to check the significance of individual Pearson’s R values (above threshold). All statistical analyses were performed using SPSS software (version 17.0; SPSS Inc, Chicago, IL). Difference was considered statistically significant at p< 0.05.

## 3. Results

### 3.1. Methanolic extract of *Tamarindus indica* inhibits HCV entry at non-cytotoxic concentration compared to other plants

Solvents with increasing polarity such as n-hexane, chloroform, methanol and water were used to prepare leaf extracts for this study. However, as several studies have suggested the beneficial effects of methanolic extracts for several plants (Table 1), we decided to initiate our study with the methanolic extract. 100µg/ ml stock solution was prepared in DMSO for each of the plants. During all the experiments in this study we used methanolic leaf extract of the four plants (*Psidium guajava L., Plumeria alba L., Syzygium cumini L., Tamarindus indica L.)* where their stock solutions were diluted with DMEM or RPMI (without FBS and antibiotics) to achieve the desired working concentration.

Cellular toxicity of all these leaf extracts was preliminarily tested on Huh7 cells by Methylene blue assay with concentration range of 10µg/ml to 1000µg/ml. From this preliminary observation, further selection was conducted by MTT assay in the given ranges: 200-700µg/ml for guava (*Psidium guajava)*, 50-300µg/ml for white champa (*Plumeria alba)*, 50-300µg/ml for black plum (*Syzygium cumini*) and 25-600µg/ml for tamarind (*Tamarindus indica)* leaf extracts. In case of *Psidium guajava* extract, we observed a significant reduction (p < 0.001) of percent cell viability at a concentration of 400µg/ml. Similarly significant reduction (p < 0.05) in cell viability was observed at 200µg/ ml for *Plumeria alba*, 75µg / ml for *Syzygium cumini* (p < 0.001) and 250µg/ ml for *Tamarindus indica* leaf extracts (p < 0.001) (Supplementary Figure 1). Optimum concentration for all the methanolic leaf extracts were selected where approximately 80% cell viability was observed and percent cell viability was not significantly reduced with their previous concentrations (Supplementary Figure 1). These optimised concentrations for each of the plant extract was selected and used in our analysis (Supplementary Table 1). Specifically, the concentrations selected were 300 μg/ml for Psidium guajava (guava); 150 μg/ml for Plumeria alba (white champa), 50 μg/ml for Syzygium cumini (black plum) and 100 μg/ml for Tamarindus indica (tamarind).

To determine the antiviral effect of these plant extracts against HCV entry in hepatocytes, Huh7 liver cells were transduced with HCVpp along with methanolic leaf extracts at their selected concentration. After 20 min of internalisation at 37°C, total RNA was isolated from transduced Huh7 cells and KGFP was estimated by qRT-PCR. Successful HCVpp entry was determined by the presence of KGFP RNA delivered within the cell. The 20 minute time point was chosen based on our previous study^11^ where we reported that by 20 minutes HCVpp and endosomal fusion has occurred and the reporter RNA (in this case KGFP) of the HCVpp had been delivered to the cytoplasm. Graphical representation of KGFP RNA detected in cells indicates that, post internalisation of HCVpp in the presence of different methanolic leaf extracts at 37°C, only the *Tamarindus indica* exhibited a significant (p< 0.05) inhibitory effect on HCVpp entry at 100µg/ml concentration (Figure 1a) in contrast to untreated control. However, *Psidium guajava*, *Plumeria alba* and *Syzygium cumini* did not show any inhibitory role at their given concentration and conditions as there was no significant difference in KGFP levels between these extracts compared to control.

**Figure 1:**
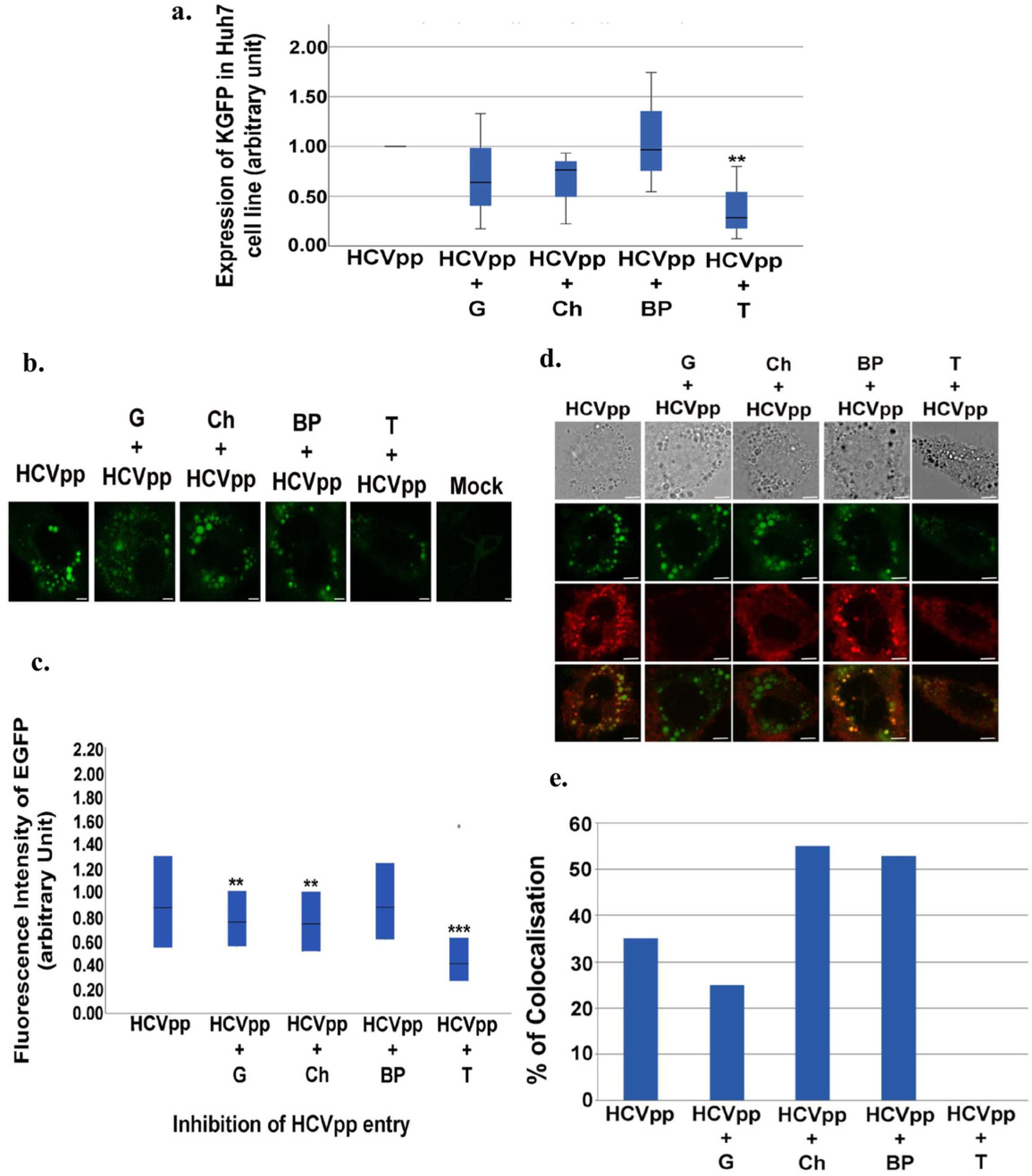
HCV entry inhibition by plant extracts. In all figures, G indicates *Psidium guajava* (guava) leaf extract, Ch: *Plumeria alba* (white champa) leaf extract, BP: *Syzygium cumini* (black plum) leaf extract and T: *Tamarindus indica* (tamarind) leaf extracts. **(a)** Graphical representation of HCVpp entry, indicated by KGFP expression in transduced Huh7 cells in the presence of plant extracts. Presence of KGFP is represented in a box plot profile. Each box represents the median, first and third quartile. Error bars indicate standard deviation, ** represents significant difference (p < 0.05) as tested by one way ANOVA followed by Post hoc Tukey (Tukey’s HSD) analysis. **(b)** Representative images of E1E2-EGFP (green) trafficking in Huh7 cells in the presence of different methanolic plant extracts. Scale Bars= 10μm. (**c**) Integrated density (IntDen) of EGFP from each of the individual infected cells is represented in a box plot profile. Each box represents that median, first and third quartile. Int Den of images for each of the experiment was counted as N. For HCVpp, N= 303; for G+ HCVpp, N= 519; for Ch+ HCVpp, N= 342; for BP+ HCVpp, N= 121; for T+ HCVpp, N= 99. ** represents significant difference (p < 0.05) and *** represents significant difference (p < 0.001) as tested by one-way ANOVA followed by Bonferonni post hoc analysis. **(d)** Representative images of E1E2-EGFP (green) trafficking in Huh7 cells in the presence of different methanolic leaf extracts colocalised with FM 4-64. Scale Bars= 10μm. (**e**) Graphical representation of positive colocalisation percentage (%) between EGFP and FM 4-64 as observed in replicate experiments. Total numbers of cells for each experiment are counted as N. For HCVPP cell lines, N= 20, N(G+ HCVPP)= 20, N(Ch+ HCVPP)= 20, N(BP+ HCVPP)= 17 and N(T+ HCVPP)= 17. t-score was determined to check the significance of Pearson’s R (above threshold) values.

### 3.2. Differential impairment at the early steps of HCV trafficking in the presence of *Plumeria alba, Psidium guajava and Tamarindus indica* leaf extracts

To confirm our previous observation to HCV entry inhibition, a parallel entry assay was conducted with fluorescent labelled HCVpp transduced Huh7 cells and their trafficking was visualised via confocal microscopy (Figure 1b). Quantification of EGFP expression in 20min post transduced Huh7 cells (Figure 1c) revealed that *Tamarindus* leaf extract treated cells (N=99) exhibited a significant difference (p< 0.001) in fluorescence expression. This was in accordance to our previous observation (Figure 1a). However, interestingly, we also found a significant difference (p<0.05) of EGFP intensity in cells treated with *Plumeria alba* (N= 342) and *Psidium guajava* (N= 519) leaf extracts, indicating reduced amount of E1E2-EGFP internalization or impaired hepatocellular trafficking. These results suggested that while delivery of EGFP labelled viral particles was similar in all cases (as per qRT-PCR results), there might be defects in HCVpp trafficking in the presence of guava and white champa leaf extracts. Consistent with our previous results black plum extracts (N= 121) did not show any significant difference in EGFP intensity compared to untreated cells.

To investigate this observation in more detail, colocalisation assay of a plasma membrane marker FM™ 4-64 with EGFP was conducted and visualised under microscope (Figure 1d). In this case, we found no colocalisation between EGFP and FM™ 4-64 in cells treated with *Tamarindus indica* leaf extract (N= 17). Methanolic leaf extract of *Plumeria alba* and *Syzygium cumini* treated cells showed higher percentage of FM™ 4-64 and EGFP co-localisation (55%, N= 20 and 53%, N= 17) compare to untreated Huh7 cells (N= 20; 35%). However, leaf extract of *Psidium guajava* treated cells showed a reduced colocalisation compared to untreated cells after 20min of viral internalization (Figure 1e). Overall, these results indicate that while *Tamarindus* extracts exhibit clear inhibitory effect against HCVpp entry, *Syzygium cumini* has no clear inhibitory role at the given concentration and conditions used in this study, although it might have an effect in viral trafficking. Extracts of *Plumeria alba* and *Psidium guajava* perhaps don’t exhibit any inhibitory effect in attachment but perhaps render some cellular trafficking defect in our experimental conditions and concentrations.

### 3.3. Selection of plant derived active compounds from *Tamarindus indica* leaf extracts to study HCV entry inhibition

We performed literature mining on *Tamarindus indica* to identify 39 compounds (Supplementary Table 2). 3D structures were obtained from PubChem and ChemSpider or drawn in MarvinSketch v20.2.0; Supplementary Table 2 lists compound identifiers and canonical SMILES, forming a curated dataset for comparative antiviral evaluation and lead prioritization.

### 3.4. Organ toxicity and toxicity endpoints analysis

SwissADME descriptors for 39 ligands (Supplementary Table 4) form two clusters: **high-lipophilicity terpenoids/sterols** (e.g., lupeol, β-sitosterol) with very high consensus Log P (≈4–9), negligible solubility, frequent Lipinski violations and moderate bioavailability—predicting formulation, dissolution and permeability challenges—and **small polar acids/sugars/vitamins** (e.g., sucrose, niacin) with high solubility but excessive H-bonding and poor passive permeation. Intermediate compounds show mixed, optimizable profiles; overall, SwissADME favors low-MW, moderately lipophilic scaffolds for oral leads and deprioritizes highly lipophilic terpenes/sterols for oral development.

### 3.5. ADME analysis

ADME predictions for 39 compounds (Supplementary Table 5) revealed two main liability clusters: **(1)** highly lipophilic sterols/terpenes and long-chain fatty acids with high LogP, poor aqueous solubility, frequent Lipinski violations, occasional hERG II flags and low predicted human tolerated doses—predicting formulation and cardiotoxicity risks despite modest LD₅₀; **(2)** small polar acids and vitamins with excellent solubility and higher tolerated doses but excessive H-bonding and poor passive permeability. A few compounds flagged AMES positivity, hepatotoxicity, or skin sensitization. Overall, small, moderately lipophilic, low-MW scaffolds are prioritized for oral development, while terpenes/sterols should be deprioritized or advanced only with structural optimization, targeted formulation, and focused in vitro ADME/toxicity follow-up.

### 3.6. Molecular Docking

To investigate whether any of the compounds are capable of interacting with HCV E2 protein, we proceeded with in silico docking. The docking results identify Lupanone, Lupeol and Triterpene as the top three in silico binders to HCV1aE2 (Table 2), with Lupanone showing the strongest predicted interaction (−8.8 kcal·mol⁻¹) followed closely by Lupeol (Figure 2a and 2b) (−8.5 kcal·mol⁻¹) and Triterpene (−7.9 kcal·mol⁻¹); when these affinities are interpreted alongside the SwissADME and PKCSM profiles, a balanced view emerges for lead selection. Lupanone combines the best docking score with very high lipophilicity and negligible aqueous solubility (consensus Log P 7.37; solubility 1.60×10⁻⁵ mg·mL⁻¹) and shows a hERG-II flag and low predicted human tolerated dose, indicating substantial formulation and cardiac-safety hurdles; Triterpene, despite a slightly weaker binding energy, has very high molecular weight and multiple H-bonding groups that predict limited passive permeability and potential metabolic instability even though it lacks obvious hERG flags. Lupeol merits particular attention for laboratory validation because it combines a strong docking score (−8.5 kcal·mol⁻¹) with a tractable safety signal: it is predicted AMES-negative, shows only moderate acute toxicity (LD₅₀ ≈ 2.56 mol·kg⁻¹), and most importantly has documented antiviral activity in the literature, making it a plausible bioactive scaffold. Its principal liabilities are very high lipophilicity (Log P 7.27), low aqueous solubility (7.69×10⁻⁵ mg·mL⁻¹) and a PKCSM-predicted hERG-II flag and low maximum tolerated dose, all of which are addressable in a staged experimental program. Accordingly, we prioritized Lupeol for immediate laboratory follow up.

**Figure 2:**
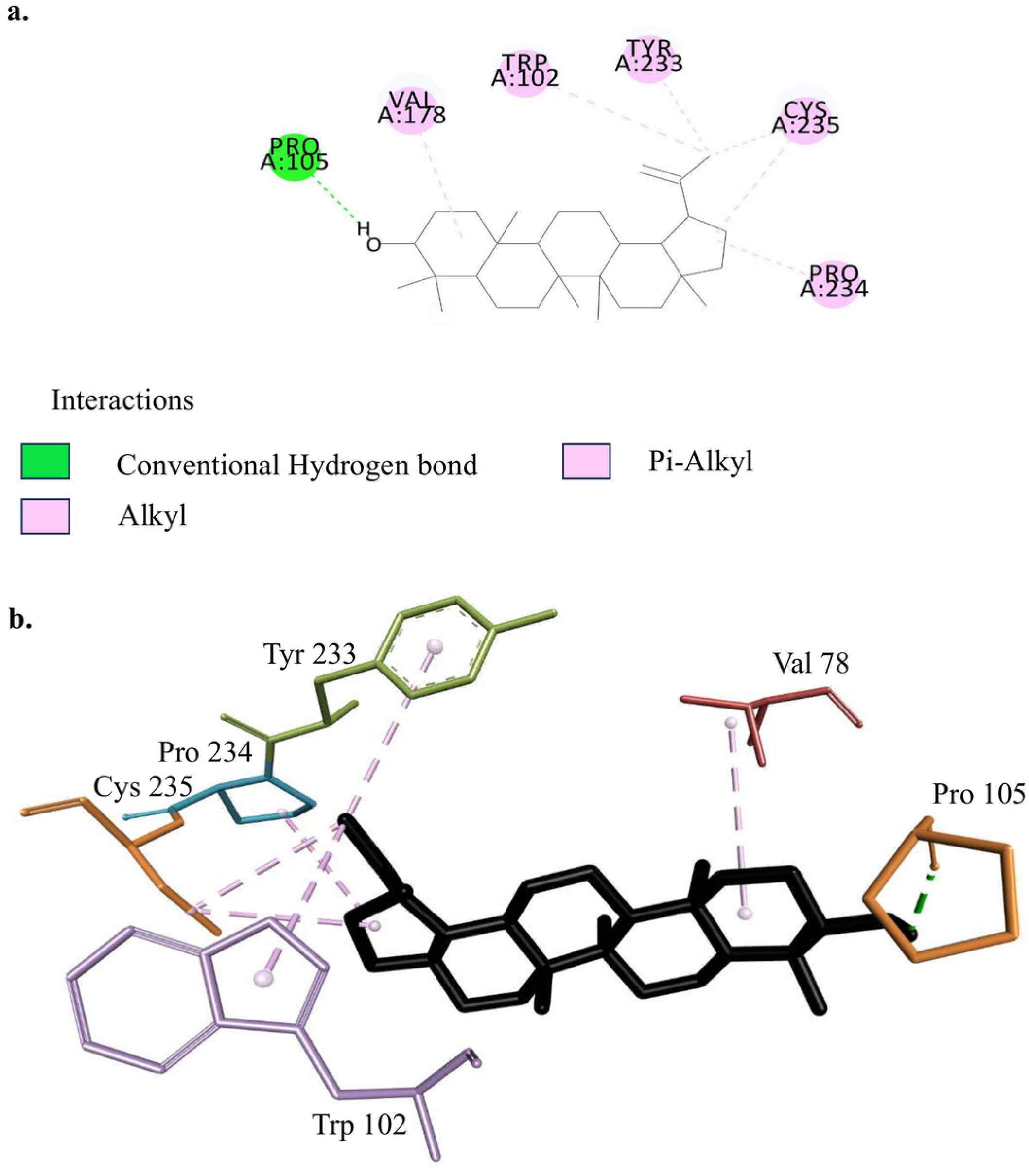
Protein-ligand interactions between Lupeol in complex with HCV1aE2. **(a)** 2D image of protein-ligand complex. The residues od HCV1aE2 that lupeol interacts with are shown. Color codes indicate the type of bond. (**b**). 3D labelled image of protein-ligand complex.

**Table 2:** Molecular docking results of antiviral compounds against HCV1aE2.

| Sl. N o. | Name of ligands | Binding Affinity (kcal/mol) | Sl. N o. | Name of ligands | Binding Affinity (kcal/mol) |
| --- | --- | --- | --- | --- | --- |
| 1 | Lupanone | -8.8 | 6 | Riboflavin | -7.4 |
| 2 | Lupeol | -8.5 | 7 | Sesquiterpenes | -7 |
| 3 | Triterpene | -7.9 | 8 | Longifolene | -6.7 |
| 4 | $\beta$_carotene | -7.8 | 9 | Caryophyllene | -6.6 |
| 5 | Cryptopine | -7.5 | 10 | Benzyl_benzoate | -6.4 |
| 11 | Linalyl_anthranilate | -6.2 | 25 | 3_eicosyne | -5 |
| 12 | 2_6_di_tert_butyl_4_methylphenol | -6.1 | 26 | Ascorbic_acid | -5 |
| 13 | B_sitosterol | -6.1 | 27 | 10_Octadecanoic_acid | -4.8 |
| 14 | Thiamine | -6 | 28 | Tartaric_acid | -4.7 |
| 15 | Sucrose | -5.9 | 29 | Niacin | -4.7 |
| 16 | Methyl_3_5_di_tert_butyl_4_hydroxybenzoate | -5.8 | 30 | Methyl_9_12_15_Octadecatrienoate | -4.6 |
| 17 | Benzene_1_2_dicarboxylic_acid | -5.6 | 31 | Malic_acid | -4.5 |
| 18 | 6_10_4_trimethylpentadeca_5_9_13_trien_2_one | -5.4 | 32 | Methyl_hexadecanoate | -4.4 |
| 19 | Pectin | -5.4 | 33 | Methyl_tricosonate | -4.4 |
| 20 | Diphenyl_ether | -5.3 | 34 | N_nona_decanoic_acid | -4.4 |
| 21 | Phytol | -5.3 | 35 | Methyl_7_10_Octadecadinoate | -4.3 |
| 22 | Hexadecanoic_acid | -5.3 | 36 | Pentadecanol | -4.2 |
| 23 | 1_Methyl_4_propylbenzene | -5.2 | 38 | Succinic_acid | -4.1 |
| 24 | Lemonene | -5.2 | 39 | Acetic_acid | -2.9 |
|  |  |  | 39 | Gum_Karaya | -1.7 |

### 3.7. Molecular Target Analysis

SwissTargetPrediction for Lupeol (Supplementary Figure 2, Supplemnetary Table 6) predicts multi-target activity: enzymes 33.3%, nuclear receptors 20%, phosphatases 20%, cytochrome P450s 13.3%, oxidoreductases 6.7%, and hydrolases/ligand-gated ion-channel targets 6.7%, indicating a preference for enzymatic and receptor-mediated interactions.

### 3.8. MD Simulation Analyses of Protein-Ligand Complexes

MD simulation (25 ns) of the Lupeol–E2 complex equilibrated within ∼5 ns and remained stable: RMSD (Figure 3a) stabilized at 0.5–0.6 nm, RMSF (Figure 3b) showed low residue mobility except for a few flexible loops, H-bonds (Supplementary Figure 3a) persisted throughout, radius of gyration (Rg) (Supplementary Figure 3b) remained ∼1.9–2.0 nm, per-residue area (Supplementary Figure 3c) values were consistent with occasional peaks at solvent-exposed residues and SASA (Supplementary Figure 3d) stayed ∼130–170 nm², and— together indicating early equilibrium and no ligand-induced backbone destabilization.

**Figure 3:**
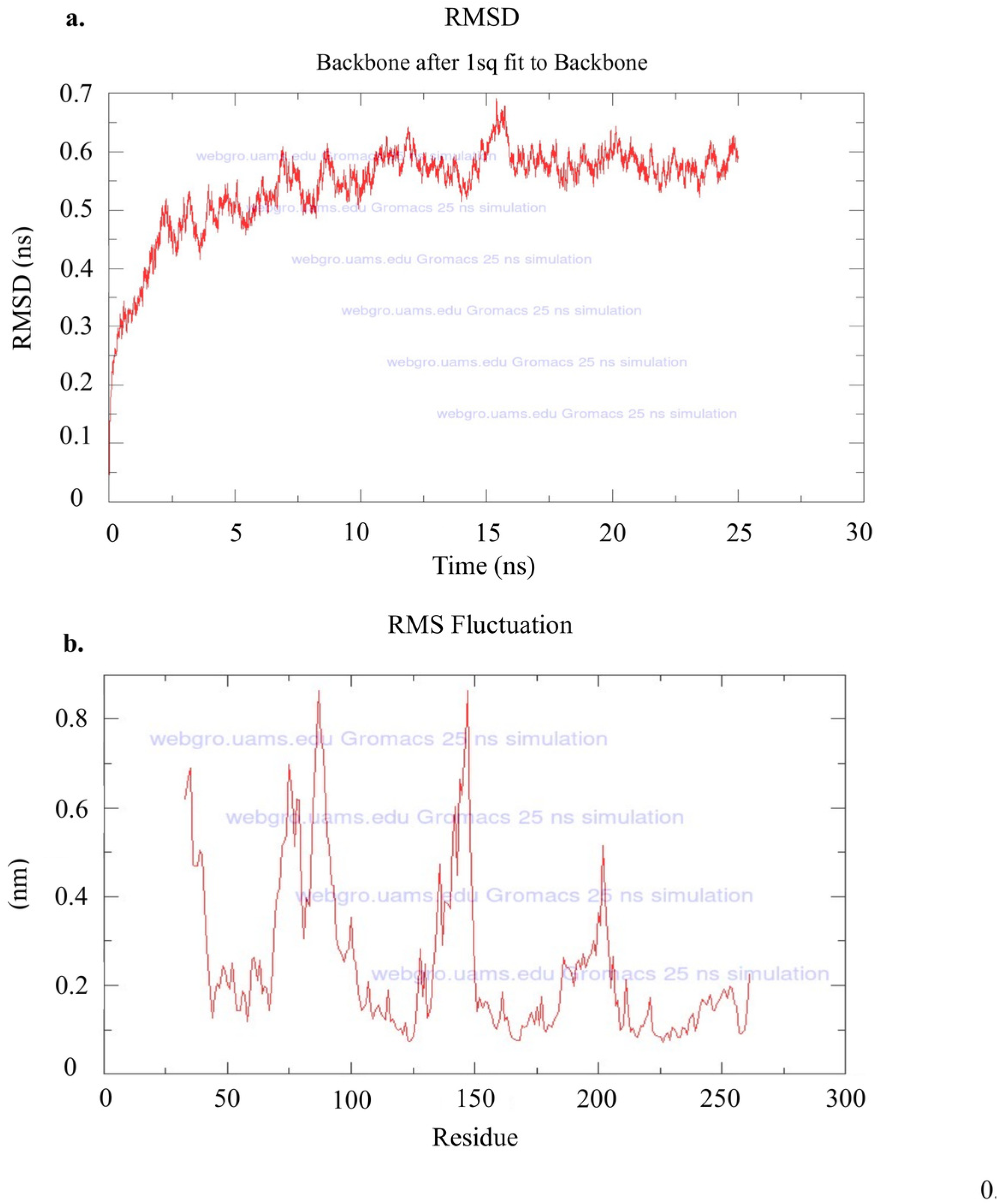
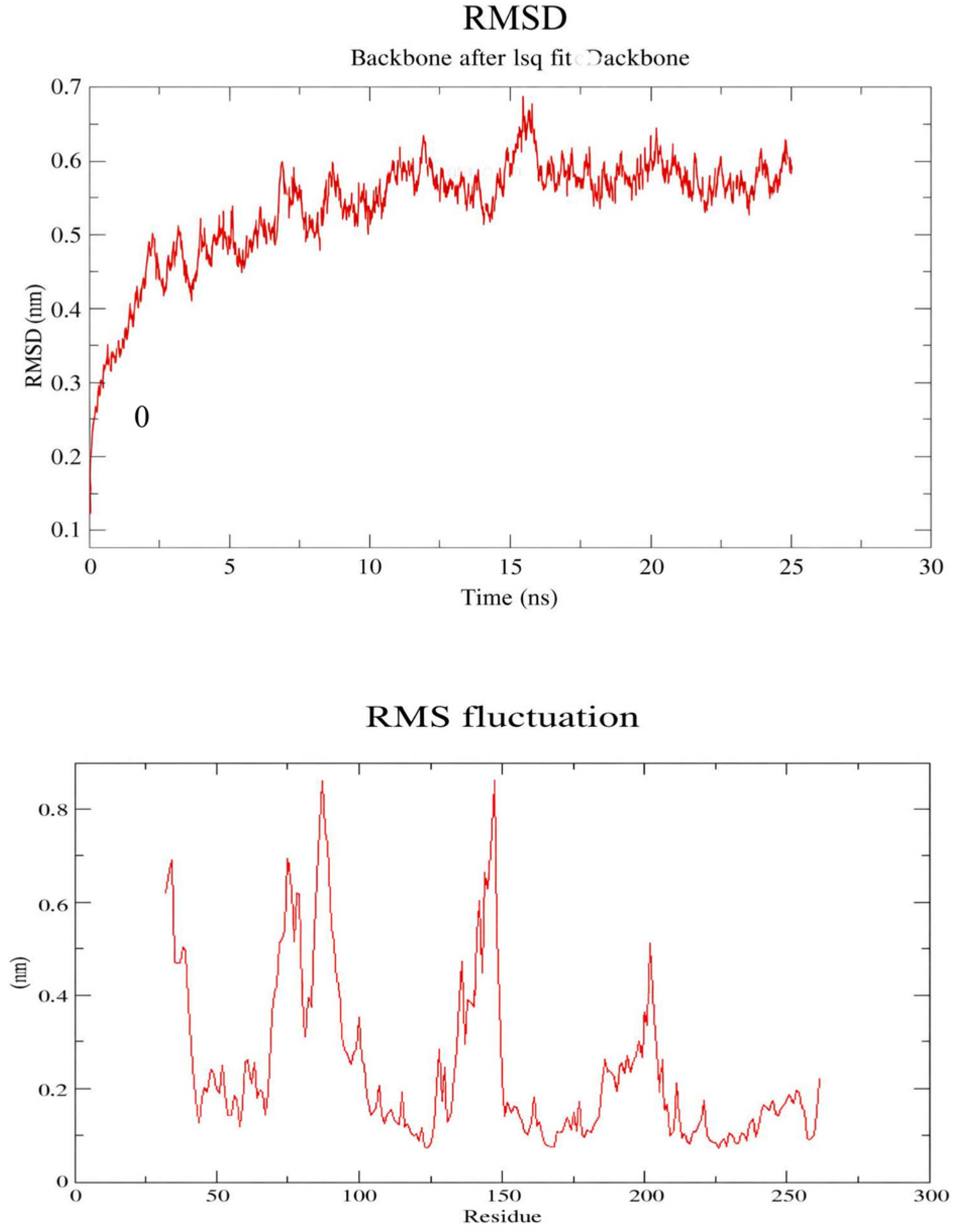
MD simulation analysis of the HCV1a E2 protein in complex with Lupeol. MD simulation was conducted for the complex for 25 ns. (**a**) Root mean square deviation (RMSD) (**b**) Root mean square fluctuations (RMSF) are depicted.

### 3.9. Transient Transfection and Binding Assay

An interaction inhibition assay was designed to study the effects of Lupeol by using the commercially available. For this experiment, a construct was used where the large extracellular domain of CD81 was expressed as a fusion with Histidine. The construct was transfected into 293 T cells and the media was collected, which had the secreted form of the CD81 LEL-His protein. CD81 LEL-His was then bound to Ni-NTA beads. A binding interaction study was then set up using the Ni-NTA beads bound to CD81-LEL-His using lysate from 293 T cells that had been previously transfected with a construct expressing E1E2 GFP. After incubation, the unbound material was saved for subsequent analysis and the beads were washed. Bound E1E2 GFP was then analysed by immunoblotting for GFP. In experiments to test the effect of Lupeol, the compound was mixed in the binding interaction mixture. It was observed that E1E2 GFP was present in the lane containing material bound to CD81-LEL-His tagged beads. This band corresponded to the E1E2 GFP band observed in the input lysate (marked as L) in Figure 4a. At the same location, there was no band in the unbound material indicating that almost all of the E1E2 GFP present in the lysate had been bound by the CD81-LEL. In the lanes containing Lupeol, the E1E2 GFP band was missing in the lane containing bound material. The E1E2 GFP band was instead visible in the lane containing unbound material. This experiment was repeated thrice, and Figure 4b depicts the densitometric estimation of the amount of E1E2 GFP bound to CD81 in the absence and presence of Lupeol considering all replicates.

**Figure 4:**
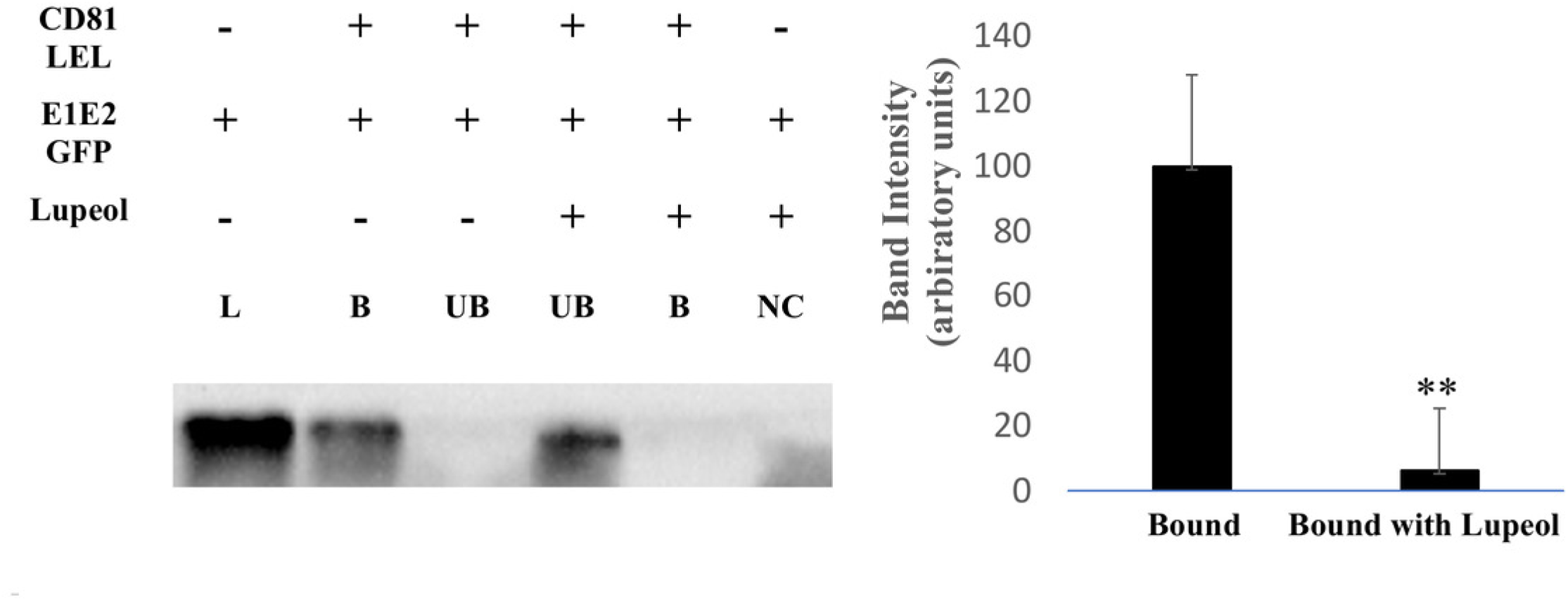
Inhibition of HCV E2 binding with CD81 in the presence of Lupeol. Co-immunoprecipitation of E1E2 GFP with CD81 LEL in the presence of Lupeol. (**a**) Western blot for E1E2 GFP is depicted. Agarose beads were pre bound to CD81 LEL. Presence and absence of Lupeol is marked as + or – as indicated. L indicates lysate, B indicates material bound to CD81 LEL containing beads and UB indicates material not bound to the beads. (**b**) Quantitation of percentage E1E2 GFP bound to CD81 LEL in the presence of Lupeol. The difference is statistically significant, *p* < 0.05 as per Student’s t -test. ** represents significant difference (p < 0.05).

Quantitation revealed that on average, 6% of E1E2 GFP bound CD81 LEL in the presence of Lupeol if the binding without the drug is considered to be 100%. This difference was seen to be statistically significant (*p* < 0.05) using Student’s t -test. This experiment proved that Lupeol was indeed capable of inhibiting the binding of E1E2 GFP to CD81-LEL.

### 3.10. Entry Inhibition Assay

To evaluate the inhibitory effect of Lupeol on HCV entry, an entry inhibition assay was performed using HCVpp. Huh7 cells were transduced with HCVpp for 15 minutes in the presence and absence of Lupeol, followed by RNA isolation and RT-PCR analysis using KGFP-specific primers. As shown in the Figure 5, lanes 1 (positive control without Lupeol) displayed distinct amplification bands at 812 nt, confirming successful viral entry. In contrast, lanes 2 (negative control without virus) showed no amplification, as expected. Importantly, lanes 3, corresponding to HCVpp infection in the presence of Lupeol, did not exhibit the 812 nt band, indicating that Lupeol effectively inhibited HCVpp entry into Huh7 cells. GAPDH expression was consistently detected across all samples, validating RNA integrity and equal loading. These findings demonstrate that Lupeol exerts a significant inhibitory effect on HCV E2-mediated viral entry.

**Figure 5:**
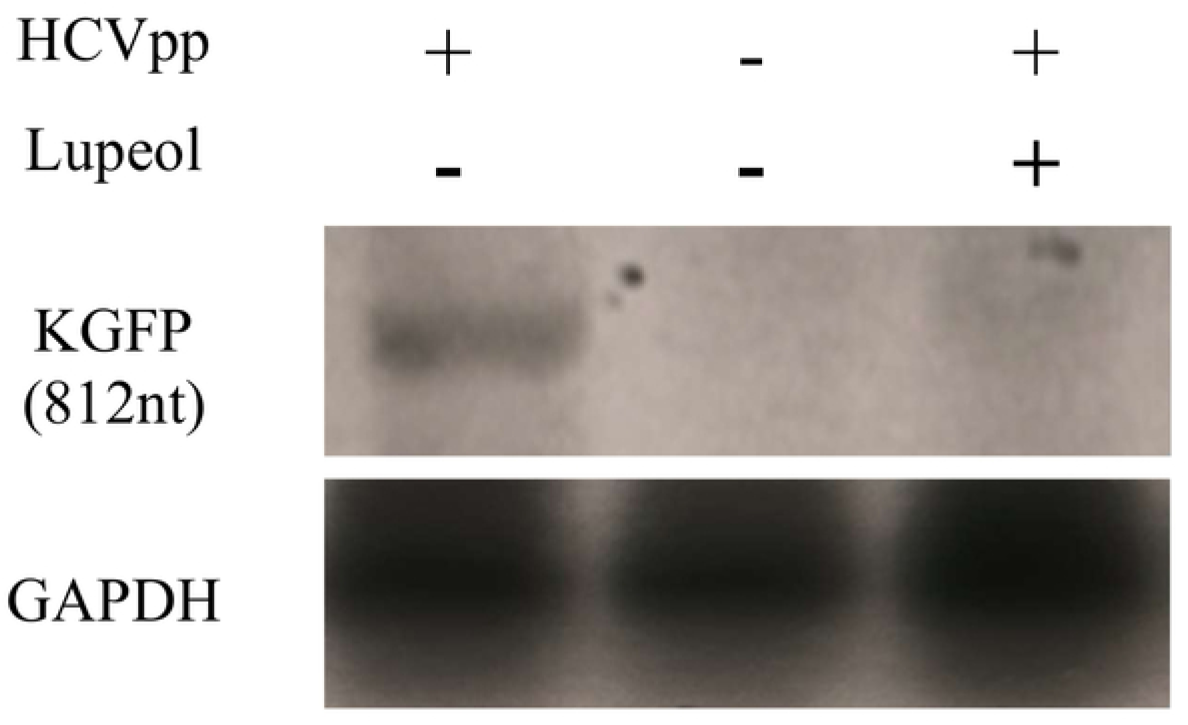
Inhibition of HCV pp entry in the presence and absence of Lupeol. Successful entry of HCVpp into Huh7 cells was detected via delivery and therefore presence of KGFP RNA (812 bp size) within the cells. Presence and absence of Lupeol are indicted as + and – respectively. Lane marked M indicates marker. Expression of GAPDH in all cell lysates are shown as loading control.

## 4. Conclusion and Discussion

The study combined ethnopharmacology, experimental virology, and computational drug discovery to identify natural HCV entry inhibitors; among four traditional Indian medicinal plants tested (*Psidium guajava, Plumeria alba, Syzygium cumini,* and *Tamarindus indica*), the methanolic leaf extract of *T. indica* showed the strongest inhibition of HCVpp entry. qRT-PCR of the KGFP reporter showed a significant reduction in viral RNA delivery with *T. indica*, and confocal microscopy of EGFP-labelled HCVpp revealed markedly reduced intracellular fluorescence, confirming impaired entry; by contrast, *P. alba* and *P. guajava* altered EGFP distribution and reduced fluorescence without significantly lowering KGFP delivery (consistent with effects on post-entry trafficking), while *S. cumini* showed no significant inhibition. The combined use of qRT-PCR, fluorescence microscopy, and plasma-membrane colocalization thus distinguished true entry blockade from trafficking perturbation and identified *Tamarindus indica* as the most promising antiviral candidate.

Having established the superior antiviral activity of *T. indica*, we next sought to identify the phytochemicals responsible for this effect. A comprehensive literature survey identified 39 phytochemicals from *T. indica* leaves and, targeting the E2–CD81 interaction to block HCV entry, molecular docking ranked Lupanone and Lupeol highest; although Lupanone scored best, Lupeol was advanced due to extensive antiviral/anti-inflammatory/hepatoprotective literature and a favorable predicted profile^22,23,24,25^. Molecular dynamics of the Lupeol–E2 complex confirmed a stable interaction, with steady RMSD, limited residue fluctuations, persistent hydrogen bonds, and minimal conformational change, supporting its candidacy for further validation.

Most importantly, the computational predictions were validated experimentally. In the biochemical interaction assay, Lupeol markedly disrupted interaction between HCV E2 and the large extracellular loop of CD81, reducing E2 binding to approximately 6% of untreated controls (p < 0.05). This finding demonstrates that Lupeol directly interferes with the receptor-binding step required for viral attachment. Consistent with this observation, Lupeol completely inhibited HCV pseudoparticle entry, as evidenced by the complete absence of KGFP amplification following pseudoparticle transduction while GAPDH expression remained unchanged. The concordance between the docking analyses, molecular dynamics simulations, biochemical interaction assay, and pseudoparticle infection assay provides strong evidence that Lupeol inhibits HCV infection by preventing E2-mediated viral entry. To the best of our knowledge, although Lupeol has previously been reported to possess antiviral activity against several RNA viruses, such as dengue virus, influenza virus, and SARS-CoV-2^26,27^, this is the first study demonstrating that Lupeol functions as an inhibitor of HCV entry through disruption of the E2-CD81 interaction.

Despite DAAs’ ^28^clinical success, HCV remains a global challenge due to cost, limited access, reinfection risk, and genotype variability^29^, and no preventive agents exist. Virus-directed entry inhibitors offer a complementary strategy to prevent initial infection or reinfection and to combine with DAAs to reduce resistance. Our findings nominate Lupeol as a promising, affordable virus-directed antiviral; given its reported activity against multiple RNA viruses, Lupeol warrants further evaluation as a potential broad-spectrum entry inhibitor.

Although the present study establishes proof-of-principle for Lupeol as an HCV entry inhibitor, several limitations should be acknowledged. The antiviral activity was evaluated using HCV pseudoparticles rather than infectious HCV, and only a single concentration of Lupeol was experimentally examined. Future studies require investigation of concentration-dependent antiviral activity, evaluate cytotoxicity of purified Lupeol over an extended concentration range, and validate these findings using infectious HCV cell culture systems and *in vivo* models. Structural studies defining the precise Lupeol-binding interface on E2, together with medicinal chemistry approaches aimed at improving solubility and pharmacokinetic properties, will further facilitate the therapeutic development of this triterpenoid.

In conclusion, this study identifies *Tamarindus indica* as a rich source of anti-HCV phytochemicals and provides the first experimental evidence that one of its major triterpenoid, Lupeol, inhibits HCV entry by disrupting the interaction between the viral glycoprotein E2 and its cellular receptor CD81. By integrating ethnopharmacological knowledge with computational screening, molecular dynamics simulations, biochemical interaction assays, and HCV pseudoparticle transduction studies, we demonstrate that Lupeol is a promising virus-directed entry inhibitor and a potential lead molecule for future anti-HCV drug development. Beyond establishing the antiviral potential of T*. indica*, this work highlights the enduring value of ethnopharmacology as a source of novel antiviral therapeutics and provides a strong foundation for developing affordable, natural-product-derived antivirals that complement existing DAA-based therapies.

## Acknowledgments

The authors acknowledge funds from DBT-BUILDER, DST-FIST, awarded to the Department of Life Sciences, Presidency University, and University Grants Commission. The authors acknowledge the expertise of Prof. Binay Chabey, Department of Botany, University of Calcutta for his help in preparation of the leaf extract. M.S. was supported by UGC.

## Conflicts of Interest

A.M. has collaboration with the Foundation for Innovations in Health (FIH) which is supported by the Common Research and Technology Development Hub in Affordable Healthcare, IIT Kharagpur; although neither FIH nor IIT KGP was involved in this present study. CB is currently employed at Premas Biotech but this organisation had no role in the work. None of the other authors have any conflicts of interest to declare.

## Declaration of the use of AI

AI tools such as Gemini was used to paraphrasing and language polishing.

---

**Supplementary Figure 1: MTT analysis for toxicity analysis of the methanolic leaf extracts.**

Cell viability in percentage depicted for (a) Psidium guajava (guava), (b) Plumeria alba (white champa), (c) Syzygium cumini (black plum) and (d) Tamarindus indica (tamarind) leaf extracts in Huh7 cells. Errors bars indicate standard deviation of three independent experiments. *** represents significant difference of p< 0.001 and ** represents significant difference of p< 0.05.

**Supplementary Figure 2: Molecular targets of Lupeol**

Predicted molecular targets of Lupeol as identified by SwissTargetPrediction analysis. The distribution of targets is shown across major macromolecular classes, including enzymes, nuclear receptors, phosphatases, cytochrome P450 family members, oxidoreductases, and hydrolases associated with ligand-gated ion channels. Percentages represent the relative proportion of each class among the predicted interactions, highlighting Lupeol’s potential for multi-target activity with a notable preference toward enzymatic and receptor-mediated pathways.

**Supplementary Figure 3: MD simulation analysis of the HCV1a E2 protein in complex with Lupeol.**

(**a**) No. of Hydrogen bonds formed between the protein binding site and the respective ligand molecules through the course of the 50 ns MD simulation. (**b**) Radius of gyration (**c**) Area per residue over trajectory. (**d**) Solvent accessible surface.

## Notes

**Funding:** This study was supported by DBT-BUILDER, DST-FIST.

